# A quantitative model for the regulation of innate immune activation

**DOI:** 10.1101/2020.04.26.061465

**Authors:** Yawei Qin, Emily M. Mace, John P. Barton

## Abstract

The immune system employs a wide variety of strategies to protect the body from infection. Cells such as natural killer (NK) cells and macrophages can recognize and eliminate targets with aberrant surface ligand expression in a manner that is not antigen-specific. This innate mechanism of activation must be tightly regulated to prevent autoimmunity. Here we introduce a quantitative model of the regulation of nonspecific activation inspired by Bayesian inference. Our model captures known behaviors of innate immune cells, including adaptation to changing environments and the development of hyposensitivity after prolonged exposure to activating signals. Our analysis also reveals a tradeoff between precision and adaptation. Maintaining the ability to adapt to different environments leads to heterogeneous responses, even for hypothetical populations of immune cells and targets that have identical surface receptor and ligand expression. Collectively, our results describe an adaptive algorithm for self/nonself discrimination that functions even in the absence of antigen restriction. The same model could also apply more broadly to the adaptive regulation of activation for other immune cell types.

## Introduction

The immune system protects the body from both internal and external threats, such as the development of cancer or infection by viruses. One common mechanism of immunity is the recognition of foreign material. T and B cells of the adaptive immune system bind specifically to antigens that are either derived from pathogens or, in the case of cancer, are altered versions of self proteins ^1^. Similarly, innate pattern recognition receptors bind to highly conserved molecules from foreign microbes, such as bacterial flagellin ^2,3^. In both cases, immune cells specifically recognize and respond to signals that do not originate from normal, healthy cells in the host.

The immune system also maintains defense mechanisms that do not rely on the specific recognition of foreign material. Immune cells like NK cells and macrophages recognize targets through a broad array of activating and inhibitory receptors, many of which bind to self-ligands. NK cells’ activating receptors primarily bind to surface ligands induced during infection, transformation, or stress ^4^. Their inhibitory receptors detect ligands that are expressed on the surface of most healthy cells, such as major histocompatibility complex (MHC) class I molecules ^5^. Similarly, many activating receptors for macrophages respond to self-ligands that indicate cellular damage, and macrophage activity is inhibited by ubiquitously expressed surface proteins such as CD47^6^.

For both NK cells and macrophages, the integration of activating and inhibitory signals is a complex, context-dependent process. NK cells were originally noted for their ability to kill target cells that express low or undetectable levels of MHC class I ^7^. However, subsequent studies showed that NK cells from mice and humans that express low levels of MHC class I are self-tolerant, though they also respond less vigorously to typical target cells ^8–12^. Patients with defects in the transporter associated to antigen processing (TAP) protein provide one such example. In these individuals, the ability of TAP to load MHC class I proteins with peptides is reduced, causing little MHC class I to reach the cell surface ^11^. Macrophages from CD47-deficient hosts are also tolerant of cells with low levels of CD47 expression, which would usually be phagocytosed ^13^. These observations point towards an adaptive process through which the threshold for activation is controlled by the local environment, which in NK cells is referred to as ‘education’ ^14,15^. This process must be extremely robust, as autoimmunity due to direct attack by NK cells and macrophages on healthy tissues is essentially unknown ^16,17^. How do these immune cells achieve self-tolerance while retaining the ability to respond to unhealthy targets? Qualitative models of education have been developed to explain NK cell behaviors ^18–20^, but no quantitative framework currently exists. Most prior quantitative work on self/nonself recognition has focused primarily on adaptive immunity and the specific recognition of nonself ^21–25^, rather than the nonspecific recognition of ‘missing’ or ‘altered’ self detected by NK cells or macrophages.

Here we use ideas from Bayesian inference to develop a quantitative model for the regulation of innate immune activation. In our model, immune cells learn the typical distribution of ligands on *healthy* cells through repeated encounters, allowing them to respond to significant deviations from typical ligand expression in rare, unhealthy targets. Our study complements mechanistic studies of immunity by focusing on the principles underlying immune regulation. The model we develop is consistent with known behaviors of NK cells and macrophages, including adaptation to different physiological environments and the development of hyposensitivity after prolonged exposure to activating stimuli. Our results also uncover a tradeoff between the precision of immune responses and the ability to adapt to different environments.

### Adaptive Regulation of Immune Activation

Immune cells respond to signals from their receptors that can be either activating or inhibitory. The strength of the signals that they receive depends on both the concentration of receptors on the immune cell surface and the concentration of the corresponding ligands on the target cell surface. We represent the net signal that an individual immune cell receives from a target cell with a single variable *x*. Larger values of *x* represent more activating signals, while smaller values represent more inhibitory signals.

To prevent autoimmunity, immune cell responses must be tuned to avoid activation against normal, healthy cells. We propose that this can be achieved by the immune cell through the construction of an internal representation *P_r_*(*x*) of the signal distribution (see **Fig. 1**). Under normal conditions, the great majority of targets that an immune cell encounters should be healthy. Signals that are substantially more activating than those from typical cells, such that *P_r_*(*x*) is very small, are likely to originate from outlier cells that may be infected, stressed, or transformed. Biologically, information needed to represent the signal distribution could be encoded by the level of intracellular proteins involved in signaling cascades, or by adjusting the distribution or spatial organization of surface receptors on the immune cell membrane.

**Fig. 1.**
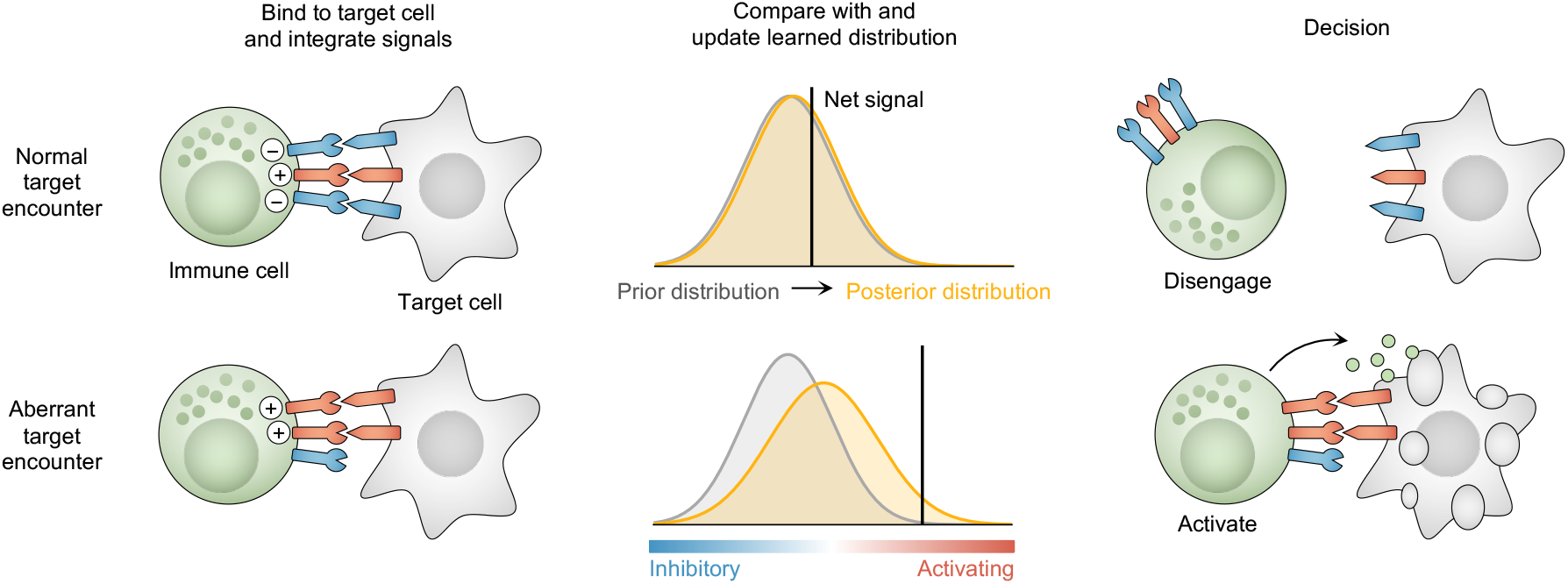
Model overview. Immune cells receive both activating and inhibitory signals from target cells that they encounter. Net signals are used to update an internal estimate of the signal distribution, reflecting the balance of activating and inhibitory ligands on target cell surfaces in the current environment. Signals from target cells that are far more activating than typical ones stimulate an immune response.

Here, we characterize the signal distribution that an immune cell receives from targets in its local environment with a Gaussian function. The true mean *μ_t_* and precision 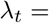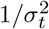 are unknown, and must be estimated through multiple encounters with target cells. Estimating the variance 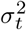 in signal values, as well as the mean, is crucial to differentiate true outlier target cells from ones with normal variation in surface ligand expression.

In general, the true signal mean and variance may be non-stationary. For example, the distribution of ligands expressed on target cells may vary as an immune cell migrates from one tissue to another. To accommodate time-varying signals, we employ a modified Bayesian update rule where the strength of the prior distribution is fixed. We begin with a normal-gamma prior for the signal mean *μ* and precision *λ*,

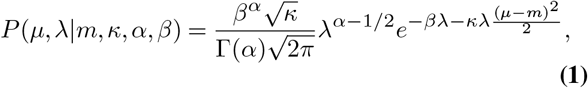

which is the conjugate prior for a Gaussian distribution with unknown mean and variance. Here Γ(*α*) represents the gamma distribution. The parameters *κ* and *α* are proportional to the number of measurements used to estimate the mean and variance, which we take to be fixed. Following equation (1), the mean value of *μ* is *m*, and the mean value of *λ* is *α/β*. When the immune cell binds to a new target cell and receives a signal *x*, the internal representation of the signal distribution is updated with new parameters

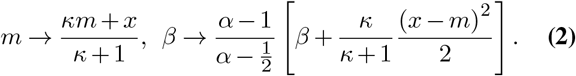

This expression differs from the standard Bayesian update in that the parameters *κ* and *α* are held fixed, and the updated value of *β* is shrunk by a factor of 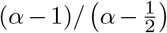 to compensate (Methods).

Our modified Bayesian inference framework introduces a *memory length* for adaptation, encoded by the parameters *κ* and *α*. The larger *κ* and *α* are, the less the estimated mean and variance will shift when a new signal is received. The update rule described in equation (2) effectively places a weight on measurements of the signal distribution that decays exponentially as new measurements are taken, emphasizing more recent signals over older ones (Methods). Larger values of *κ* and *α* result in a slower decay, and thus the memory of the signal distribution becomes longer. We emphasize that this notion of memory is distinct from the concept of memory in adaptive immunity. In this context, memory introduces a tradeoff between precision and adaptability. When the memory is long, it is possible to adapt to a specific distribution precisely, but adaptation to new environments is slow.

Integrating over the unknown mean and precision in equation (1), the internal representation of the signal distribution *P_r_*(*x*) takes the form of a shifted, scaled t-distribution (Methods). During each target cell encounter, we the immune cell activates if it receives a signal *x* that is substantially more activating than typical signals from the estimated distribution.

Specifically, we assume that activation occurs when

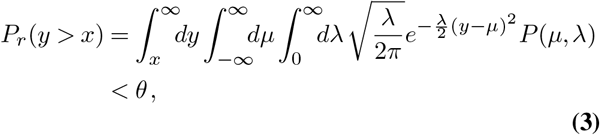

for some threshold value *θ*. The threshold *θ* must be small to avoid autoimmunity. If an immune cell were to learn the exact signal distribution in an environment where all target cells are healthy, then *θ* would be the probability that the immune cell activates against a healthy cell. Here we will treat *θ* as constant, but in principle *θ* could be modulated adaptively by the immune system, for example to provide heightened surveillance during infection.

We propose that some behaviors of immune cells like as NK cells and macrophages can be understood by considering how they adapt to typical levels of stimulation from target cells in the local environment. As we show in Results, this perspective is consistent with past experimental results. Our model also highlights the inherent tradeoff between precision and the ability to adapt to different environments.

## Results

### Immune cells adapt to the signal distribution in a static environment

We first tested the ability of our model to recover the true parameters of a test signal distribution. We compared the estimated values of *μ* and 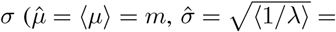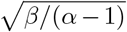, where 〈·〉 denotes an average over equation (1)) to those of the true signal distribution. **Figure 2** shows that immune cells adapt to the distribution of signals in the environment, even when the initial parameters of the prior are far from the true ones. The number of target cell encounters required to approach the true parameters depends on the initial distance from them, and on the memory length. The shorter the memory length, the faster the convergence (**Supplementary Fig. 1**).

**Fig. 2.**
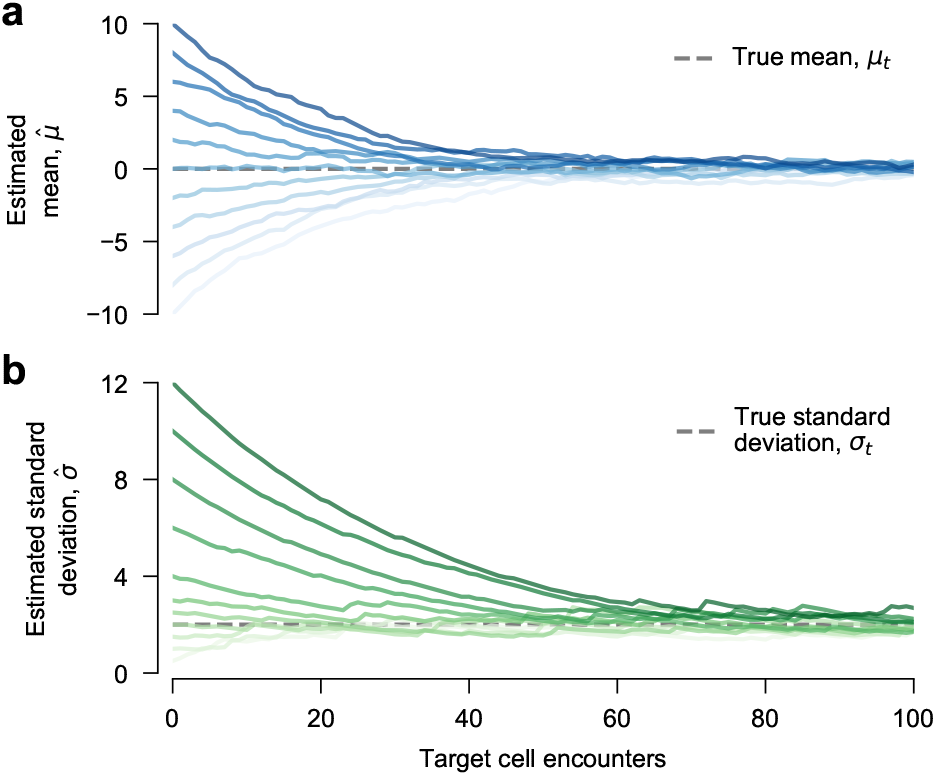
Immune cells adapt to a static environment. The mean 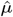 and standard deviation 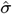 of the internal representation of the signal distribution converge toward the true mean (*μ_t_* = 0) and standard deviation (*σ_t_* = 2) of the signal distribution in the environment. **a**, Convergence to *μ_t_* from various initial values of *m*. The initial value of *β* = (*α* − 1) is the same in all cases. **b**, Convergence to *σ_t_* from various initial values of *β*. The initial value of *m* = 0 is the same in all cases. For both sections, the memory length is set by *κ* = 20 and *α* = 10.

Adaptation to the true signal mean and variance is not perfect, however. This is because the memory length is finite. Shorter memory lengths result in greater ‘noise’ in the inferred parameters of the signal distribution.

### Immune cells adapt to the signal distribution in dynamic environments

NK cells show a remarkable ability to adapt to changing environments. As noted above, MHC class I is a powerful inhibitory ligand for NK cells. NK cells in hosts that naturally express low levels of MHC class I are self-tolerant. Yet, they are also hyporesponsive to MHC class I-deficient target cells that would usually be killed by NK cells from hosts with normal levels of MHC class I expression ^8,10^. A pair of experiments showed that these behaviors are dynamic, rather than being fixed during development. When mature NK cells were transferred from MHC class I-deficient mice into mice with normal MHC class I expression, they regained their ability to kill MHC-deficient targets ^26,27^. Conversely, NK cells from normal mice that were transferred into the MHC class Ideficient environment became hyposensitive ^26^. Importantly, this shift in behavior occurs without the need for cell division or changes in the composition of receptors on the NK cell surface ^26^. Additional experiments have confirmed that the manipulation of MHC class I expression in mice leads to analogous results ^28,29^.

Adaptation also occurs when NK cells or macrophages undergo prolonged exposure to activating stimuli. For macrophages, extended exposure to lipopolysaccharide (LPS) results in blunted responses to additional stimulation by LPS, a phenomenon known as endotoxin tolerance ^30,31^. When LPS is withdrawn, macrophages gradually recover normal function ^30^. In NK cells, a similar phenomenon has been observed with prolonged exposure to ligands for the activating receptor NKG2D, resulting in hyposensitivity ^32–34^. NK cells desensitized through long-duration exposure to NKG2D ligands were also hyposensitive to other activating stimuli ^33^. Similar results have also been observed for prolonged exposure to ligands for other activating receptors ^35–37^.

We ran a series to simulations to mimic the transient exposure of immune cells to different environments or levels of stimulus. In our simulations, immune cells began in a ‘normal’ environment with *μ_t_* = 0 and *σ_t_* = 1. To quantify their ability to activate against targets that express high levels of activating ligands and/or low levels of inhibitory ligands, we computed the probability that an individual immune cell activates in response to an aberrant target cell with a signal drawn from a Gaussian distribution with mean *μ_a_* = 5 and standard deviation *σ_a_* = 2, using a threshold *θ* = 0.01. Because *μ_a_* is substantially higher than *μ_t_*, due to the presence (lack) of activating (inhibitory) ligands on the target cell surface, the probability of activation is initially high (**Fig. 3**).

**Fig. 3.**
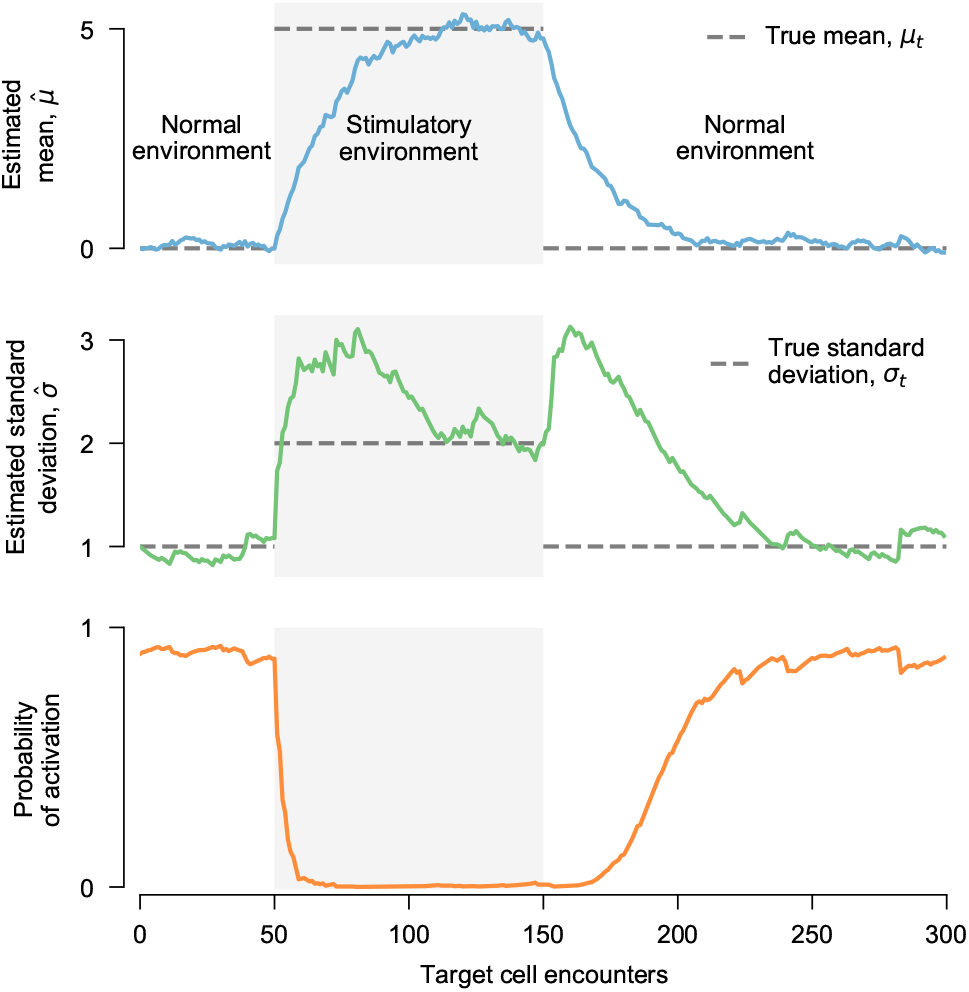
Immune cells adapt to changing environments, mimicking experimentally observed development of hyposensitivity and recovery. Adaptation of the estimated signal mean 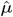, standard deviation 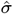, and probability of activation against an aberrant target as an immune cell is transferred between different environments. In the initial, ‘normal’ environment, the level of activating stimulus is low and the immune cell is primed to respond to aberrant targets. After transfer to a new, more stimulating environment (shaded region), the immune cell adapts to the new signal distribution and progressively loses the ability to respond to aberrant targets. When the immune cell is returned to the ‘normal’ environment, responsiveness is gradually restored. To compute the probability of activation, we used a Gaussian signal distribution from aberrant targets with mean 5 and standard deviation 2, which matches the environment in the shaded region. Initial parameters of the estimated signal distribution are *m* = 0 and *β* = 9 (leading to 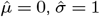), with memory parameters *κ* = 20 and *α* = 10.

To simulate a change in environment, after 50 encounters with normal targets we switched the distribution of signals to match the signal distribution from ‘aberrant’ targets described above. As the immune cell adapts to this new environment, its probability of activation by aberrant target cells decreases. The probability of activation settles near zero once the immune cell fully adapts to the environment. However, this hyposensitivity is not permanent. After a total of 150 target cell encounters, we return the immune cell to the normal environment (*μ_t_* = 0, *σ_t_* = 1). As the immune cell encounters more healthy cells, it regains its capacity to activate against aberrant targets. This trajectory mimics the development of hyposensitivity and restoration of normal function described in experiments ^26,27,30–37^, and is observed consistently in our simulations (**Supplementary Fig. 2**).

The number of encounters required to develop hyposensitivity in **Fig. 3**, and to recover it after the immune cell returns to the normal environment, depends on the memory parameters *κ* and *α*. This behavior also depends on the difference between the normal and aberrant environments. However, the development and loss of hyposensitivity is a robust prediction of our model that does not depend on a precise choice of parameters. Biologically, the memory length could be adaptively modulated by the immune system, for example through cytokine signaling, to better protect against infection. We also considered a modified model where signals were sigmoidally transformed, mimicking a saturation effect for strong activating or inhibitory stimuli (Methods). This model makes the process of adapting to different environments more gradual (**Supplementary Figs. 3-4**).

### Finite memory results in heterogeneous immune cell responses

As shown in **Fig. 2**, immune cells in our model do not adapt perfectly to the true signal distribution in the environment due to finite memory. Together with the stochastic nature of target cell encounters, this implies that there will be a range of immune cell responses, even among immune cells with identical receptors. **Supplementary Figure 2** shows an example of this behavior in a population of immune cells transferred between different environments, as shown for a single cell in **Fig. 3**.

To systematically explore this heterogeneity, we sought to characterize the distribution of estimated signal parameters *m* and *β* for a population of identical immune cells with finite memory, which inhabit the same environment. As an analytical approach, we developed a continuous approximation of the discrete modified Bayesian update dynamics described in equation (2). Assuming that parameter updates with each target cell encounter are small, equation (2) is described by a set of stochastic differential equations (Methods). We then derived a Fokker-Planck equation that describes the evolution of the distribution of learned *m* and *β* values for a population of immune cells with identical receptors in the same environment (Methods).

**Figure 4a** shows one example of the learned distribution of *m* and *β* parameters, derived from numerical integration of the Fokker-Planck equation. As expected, the distribution of learned parameters is centered around the true ones. Due to finite memory, some immune cells that have recently interacted with targets that provide unusually low levels of stimulus underestimate the true signal mean in this environment, for example.

**Fig. 4.**
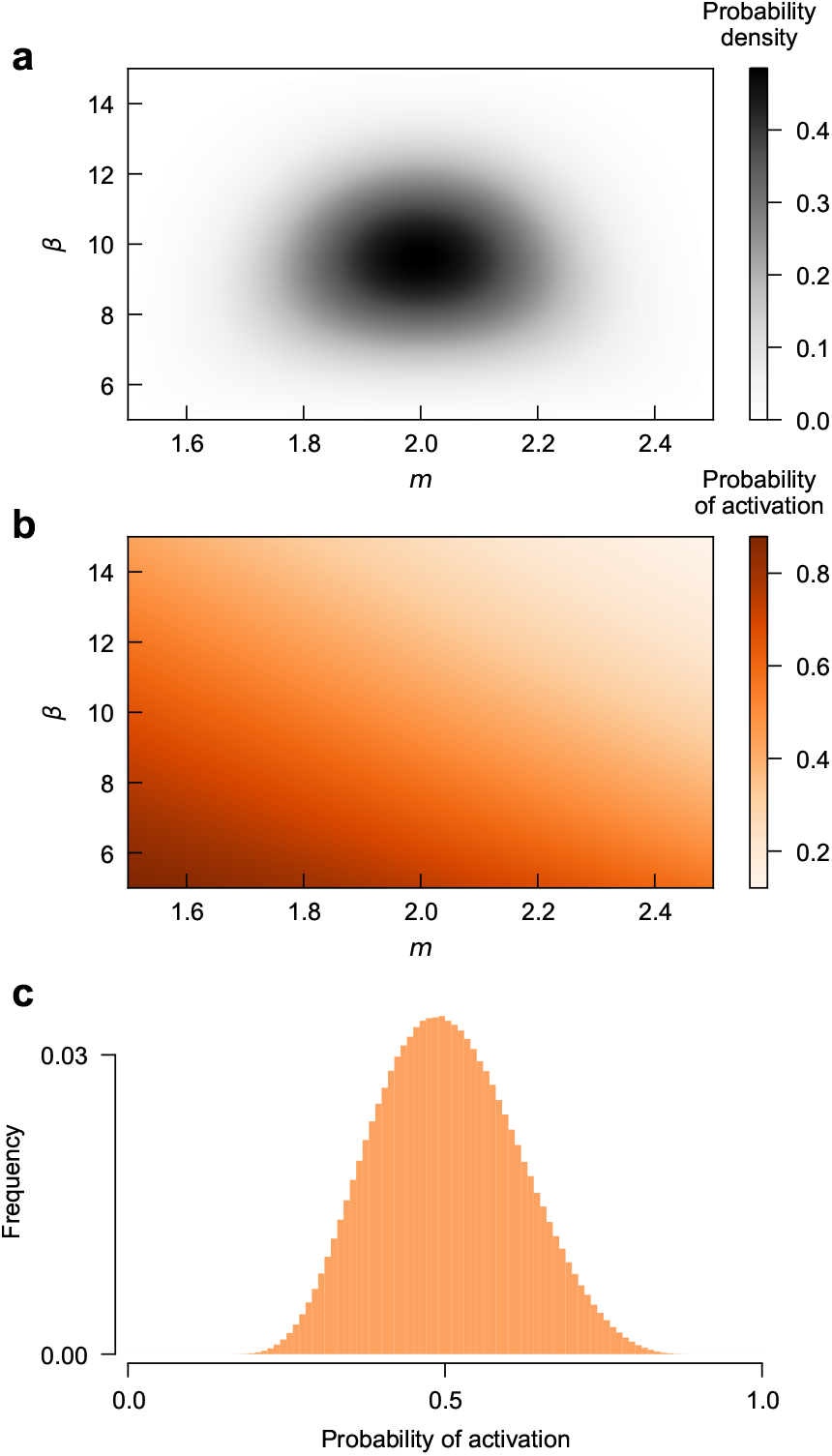
Steady state distribution of immune cell adaptation and responsiveness due to finite memory. **a**, Steady state joint probability distribution of learned (*m, β*) parameters, estimated by numerical solution of the Fokker-Planck equation. Here the true signal mean is *μ_t_* = 2, and the true standard deviation is *σ_t_* = 1. Here we used memory values *κ* = 20 and *α* = 10. We observe that the learned parameters are indeed concentrated around the true ones 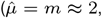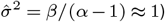. **b**, Probability of activation against aberrant targets with signal distribution *μ_a_* = 4.5 and *σ_a_* = 1 as a function of (*m, β*). Immune cells that happen to have lower values of both *m* and *β* have higher confidence that target cell signals should be more inhibitory, and thus they are more sensitive to stimulus from aberrant targets. **c**, Net distribution of probabilities of activation against aberrant targets, obtained by combining the distributions in **a** and **b**. A wide range of responses exist: some immune cells have a high probability of recognizing aberrant targets, while others are relatively unlikely to respond.

Heterogeneous adaptation to the environment results in heterogeneous responses to target cells. **Figure 4b** shows the probability that an immune cell with particular values of *m* and *β* responds to a set of aberrant target cells. Lower values of *m* and *β* result in estimated signal distributions *P_r_*(*x*) that are more concentrated around smaller values of the signal *x*, making these immune cells more sensitive to aberrant targets (see equation (3)). Thus, even for immune cells that express identical sets of receptors, stochastic encounters lead to a range in responsiveness to targets (**Fig. 4c**).

Memory length controls the degree of heterogeneity in immune cell adaptation to the environment, and consequently, the degree of heterogeneity in immune cell responses to targets. **Figure 5** shows responses of a panel of immune cell populations with different values of the memory parameters *κ* and *α*. Shorter memory values result in more heterogeneous responses. Responses against aberrant targets are more reliable for cells with longer memories. But importantly, even fairly short memory lengths result in behavior that is qualitatively similar to immune cells with longer memories and greater precision. Short memory lengths do not necessarily lead to pathological responses such as increased activation against healthy targets or hyposensitivity to strongly activating target cells. And as noted previously, long memories result in slow adaptation to different environments (**Supplementary Fig. 1**).

**Fig. 5.**
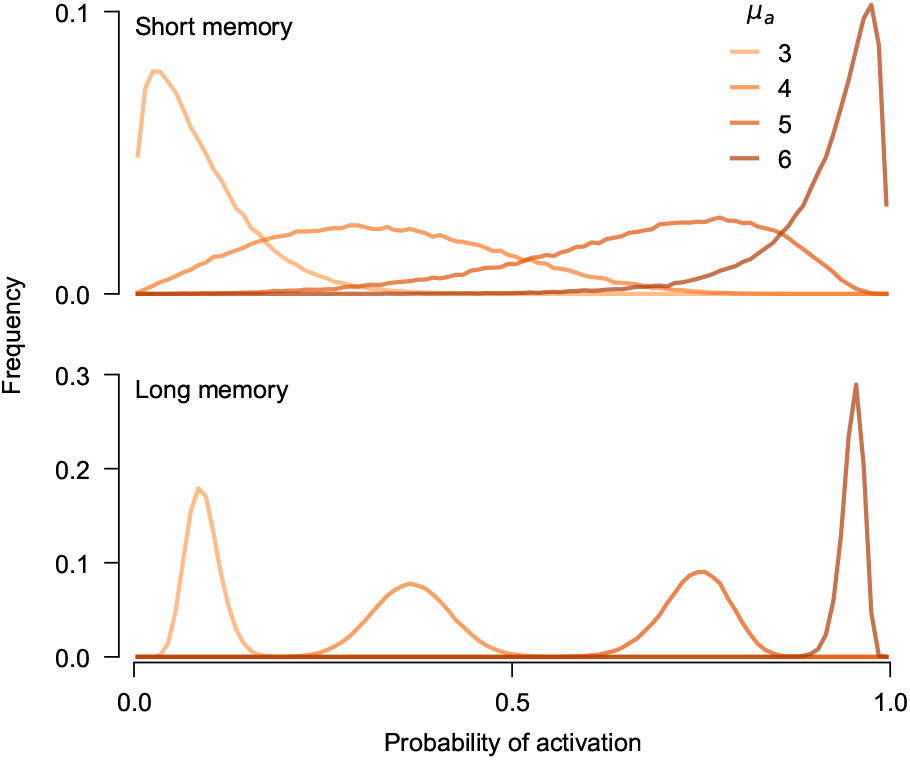
Immune cell responses are diverse, but follow predictable trends depending on the level of stimulation received from target cells. Distribution of probabilities of activation for populations of immune cells with short (*κ* = 10, *α* = 5, *top*) and long (*κ* = 100, *α* = 50, *bottom*) memory lengths. Each immune cell population consisted of 10^5^ cells with random initial values of *m* and *β* uniformly distributed between [0, 5] and [0, 40], respectively. Immune populations were evolved through 10^3^ target cell encounters with signal distributions *μ_t_* = 2 and *σ_t_* = 1. Immune cells were then tested for their probability to activate against aberrant targets that provide different levels of stimulus (*μ_a_* = (3, 4, 5, 6)). Here *σ_a_* = 1 in all cases. For both shorter and longer memory lengths, the probability of activation is low when *μ_a_* is close to *μ_t_* and high when *μ_a_* ⨠ *μ_t_*. However, the spread in activation probabilities is significantly larger for immune cell populations with short memories.

Our results on immune cell heterogeneity are consistent with experimental studies. Investigations of the patterns of target cell killing have observed widely varying responses for individual NK cells: some kill many targets efficiently, while others are inactive ^38–43^. This finding holds not only for diverse primary cells ^38–40^, but also for NK cell lines ^41–43^, where cells in the population would be predicted to have homogeneous expression and density of receptors.

## Discussion

Theoretical analyses of immunity have often focused on the adaptive immune system and antigen-specific recognition of foreign material to distinguish self from nonself ^22,44–46^. Here, we described an ‘algorithm’ for self/nonself discrimination that operates in a very different way. Rather than learning to detect specific pathogens, our model immune cells learn the properties of *healthy* cells in their current environment, which allows them to respond to aberrant cells that may be infected, stressed, or transformed. Learning in our model is ‘unsupervised’ in the sense that immune cells do not have external information about whether the targets that they encounter are healthy or not. Nonetheless, it operates reliably following the simple assumption that the great majority of targets that immune cells encounter are likely to be normal, healthy cells.

Our model captures multiple experimentally observed behaviors of innate immune cells. These include adapting responses to different environments, the development of hyposensitivity after prolonged exposure to stimulus, and the eventual recovery of normal function after the stimulus is withdrawn. At present, however, little data exists to quantitatively test predictions for how past encounters with target cells affect future responses. Future measurements of the kinetics of adaptation would be of great interest.

Recent work has also applied ideas Bayesian inference to immunity, focusing in particular on rules for optimizing the adaptive immune repertoire ^46,47^. There, models were constructed to optimally allocate immune cells with different antigen specificities to combat a shifting environment of pathogens. Our work is similar in that we also consider adaptation to varying environments. However, our model concerns the adaptation of single immune cells, and we make no assumptions about how the local environment will vary over time. Another intriguing study developed a machine learning-based model of negative selection in T cells, where encounters with self peptides are central ^48^. More generally, the problem of estimating time-varying signal distributions has some similarities with estimation using Kalman filters ^49,50^. An important difference in the present case is that the signal variance must also be estimated, and the way that the signal mean and variance change is unknown.

Similarly, our model can be compared with the discontinuity theory of immunity ^51^, which posits that immune cells in general respond to sharp changes in the environment. One of the main differences between our model and the discontinuity theory is that we explicitly consider the *variance* of signals in the environment. Though additional experiments will be needed to explore these models in quantitative detail, there is some evidence that variance in ligand expression is important. A recent experiment showed that MHC class I-deficient hematopoietic cells are spared in mice that also have hematopoietic cells with normal levels of MHC class I expression, but only if the MHC class I-deficient cells comprise a substantial fraction of all cells ^29^.

Our model can be extended to incorporate additional features of immune-target interactions. One important extension of the model would be to explicitly consider signaling through multiple pairs of receptors and ligands. NK cells and macrophages use a wide array of receptors ^5,6^. Intriguingly, recent studies have revealed dramatic heterogeneity in the complement of receptors that individual NK cells express ^52,53^. Quantifying the ability of populations of immune cells with different patterns of receptor expression to collectively recognize target cells could shed light on principles governing the innate immune repertoire.

Though we have focused on cells of the innate immune system, inspired especially by NK cells and macrophages, the model we developed may also apply more broadly to other cell types. There are some conceptual similarities between our model and the ‘tunable threshold’ model, originally applied to T cell signaling ^54,55^. Recent work demonstrated that T cells adapt to basal levels of T cell receptor (TCR) signaling, and that cells with stronger basal signaling were less responsive ^56^. Importantly, this work also demonstrated that even T cells with identical TCRs exhibit heterogeneous responses to stimulus ^56^, which is one of the main predictions of our model.

## ACKNOWLEDGEMENTS

Computations were performed using the computer clusters and data storage resources of the UCR HPCC, which were funded by grants from NSF (MRI-1429826) and NIH (1S10OD016290-01A1). This work is supported in part by R01AI137073 to EMM.

## AUTHOR CONTRIBUTIONS

Author contributions: Y.Q. and J.P.B. designed research; Y.Q. and J.P.B. performed research; Y.Q., E.M.M., and J.P.B. analyzed data; Y.Q., E.M.M., and J.P.B. wrote the paper.

## Methods

### Inference methods

#### Bayesian inference for a Gaussian distribution with unknown mean and precision

Given a normal-gamma prior distribution for the Gaussian mean *μ* and precision *λ*, the posterior distribution for *μ* and *λ* after taking a new measurement *x* is

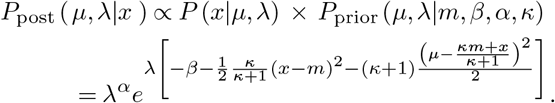

Recall that

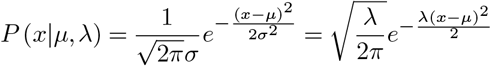

is the likelihood function, where *λ* = 1*/σ*^2^ is the precision, and

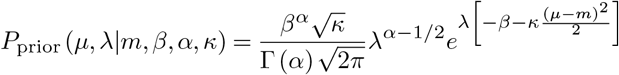

is the normal-gamma prior. The posterior thus also follows a normal-gamma distribution with modified parameters *m*′, *β*′, *α*′, and *κ*′, which are related to those of the prior distribution by

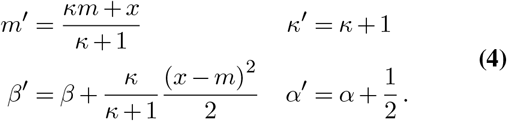

#### Estimating the signal mean and variance

The posterior signal mean 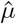 and variance 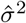 are obtained by averaging over the posterior distribution

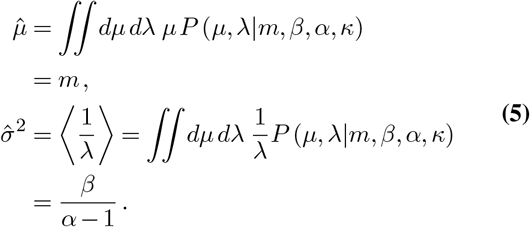

After receiving another signal *x*, the new posterior mean and variance become

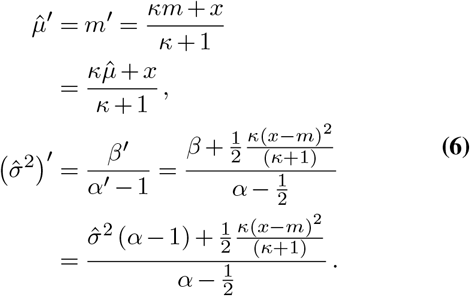

#### Reconstructed signal distribution

In our model, the internal representation of the signal distribution *P_r_*(*x*) is calculated by integrating over Gaussian distributions with mean *μ* and precision *λ*, weighted by the normal-gamma prior *P* (*μ, λ|m, κ, α, β*). Performing this integration gives

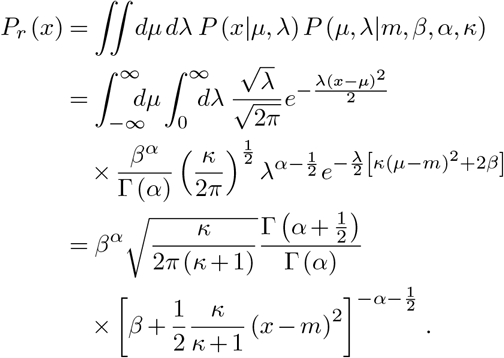

This distribution above is a scaled, shifted t-distribution

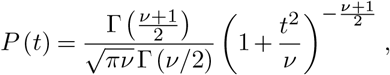

where 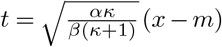 and *ν* = 2*α*.

#### Modified inference to adapt to multiple environments

While the standard Bayesian approach described above works well for static distributions, it is not appropriate for a distribution that changes with time. To enable adaptation to dynamic environments, we modify the update rules by fixing *κ* and *α*, which determine the influence of each successive measurement. Effectively, these parameters count the number of samples used to estimate the mean and precision, respectively. To ensure that estimates of the variance remain finite while *κ* and *α* are fixed, *β* must be rescaled with each update

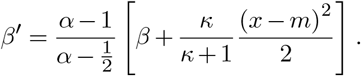

This not only ensures that *β* is finite, it also gives the same expression for updating the estimated variance as above,

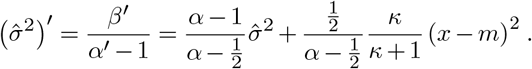

In summary, the adaptive update rules are given by

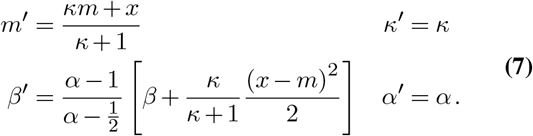

#### Modified inference as exponential weight decay

In standard Bayesian inference, all measurements contribute equally to parameter estimation. Let {*x*_1_*, x*_2_*,…,x_n_*} represent a set of *n* signals, where the subscript indicates the order in which each signal was measured. Assuming starting prior parameter values of *m*_0_, *β*_0_, *α*_0_ and *κ*_0_, and following the standard update rules given in equation (4), after *n* signal measurements we have

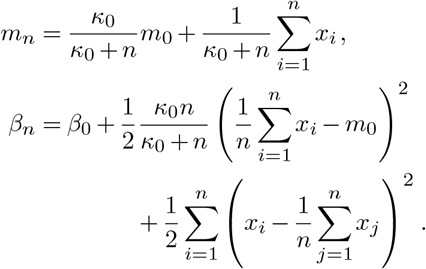

If we instead follow the adaptive update rules given in equation (7), the parameter estimates become

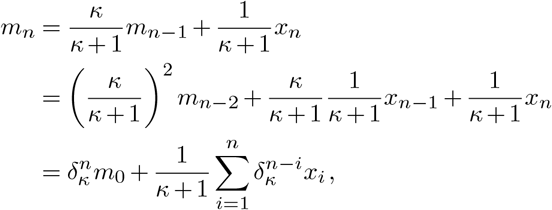

where *γ_κ_* = *κ*/(*κ* + 1), and

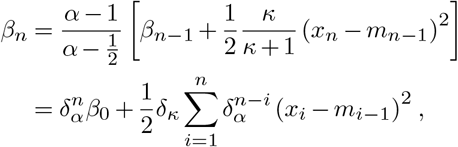

where *δ_α_* = (*α* − 1) / (*α* − 1/2). Here, each of the *n* measurements no longer contributes equally to the estimated parameters. New measurements are emphasized more strongly than old ones. With each successive measurement, the effective weight of older signal measurements decreases by a factor of *δ_κ_* for *m*, and by a factor of *δ_α_* for *β*.

The argument above facilitates the interpretation of *κ* and *α* as parameters controlling the memory length for estimating the signal distribution. Though it would be natural to take *κ* ~ 2*α*, in principle the memory length for *m* and *β* can be separately controlled by adjusting *κ* and *α*, respectively. Larger values of *κ* and *α* result in longer memory lengths and slower adaptation.

#### Inference with signal saturation

In the analysis above, the range of the signal *x* is in principle unbounded. However, immune cells have a finite number of receptors, limiting the magnitude of activating or inhibitory stimulus that they can receive. To mimic this saturation effect, we considered sigmoidally transforming signals *x* relative to the current estimated signal mean *m*. Specifically, we considered a transformed signal *x*′ given by

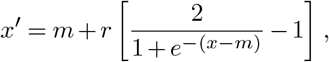

where *r* sets the range of signals that can be received, which lies in [*m* − *r*, *m* + *r*]. Because strong activating and inhibitory signals are clipped to a finite range, adaptation to environments that differ substantially from the current one is slower (see Supplementary Figs. 3-4). In this model, the estimated mean ultimately converges to the true signal mean but the standard deviation converges to a different, smaller value (which we call the transformed standard deviation, *σ_tf_*) because the sigmoidal transformation reduces the signal variance.

### Fokker-Planck equation

#### Derivation of the stochastic differential equations

Here we attempt to characterize the evolution of the estimated signal distribution parameters across a large population of cells with identical receptors. To do this we begin by using the parameter update rules, given in equation (7), to take the difference between the new the updated parameters,

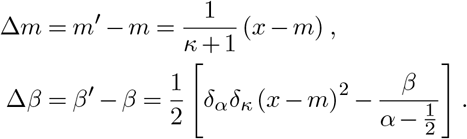

For an immune cell that is well-adapted to the environment, Δ*m* and Δ*β* should be small, especially when the memory parameters *κ* and *α* are large. Writing the time interval Δ*t* as a single encounter, we can develop stochastic differential equations that describe the evolution of *m* and *β*,

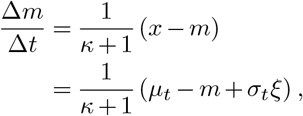

and

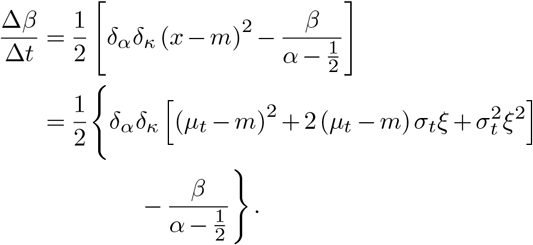

Here we assume a constant environment, where the signal distribution is Gaussian with mean *μ_t_* and standard deviation *σ_t_*. We thus write *x* = *μ_t_* + *σ_t_ξ*, where *ξ* is a Gaussian white noise.

Collectively, the evolution of ***θ*** = (*m, β*) follows

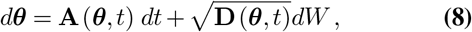

where **A** and **D** are referred to as the drift vector and the diffusion matrix, respectively. The drift vector describes the expected change in ***θ*** parameters, and the diffusion matrix describes the covariance of changes in ***θ***,

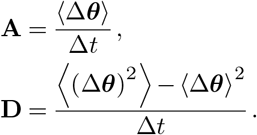

This allows us to write down the cumulants

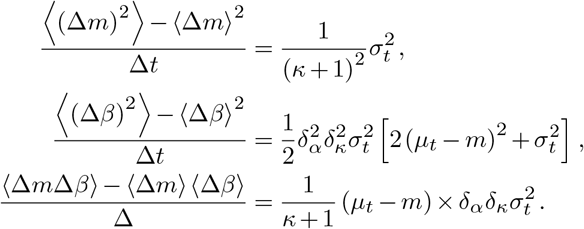

Finally we obtain expressions for the vector **A** and matrix **D**,

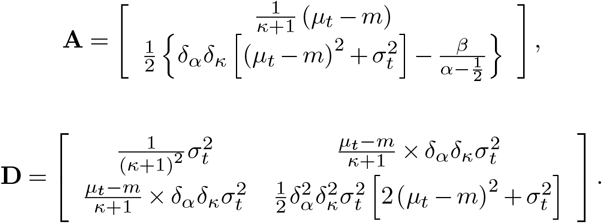

The components of **A** and **D** can thus be computed by finding the first and second moments of Δ***θ***, which are

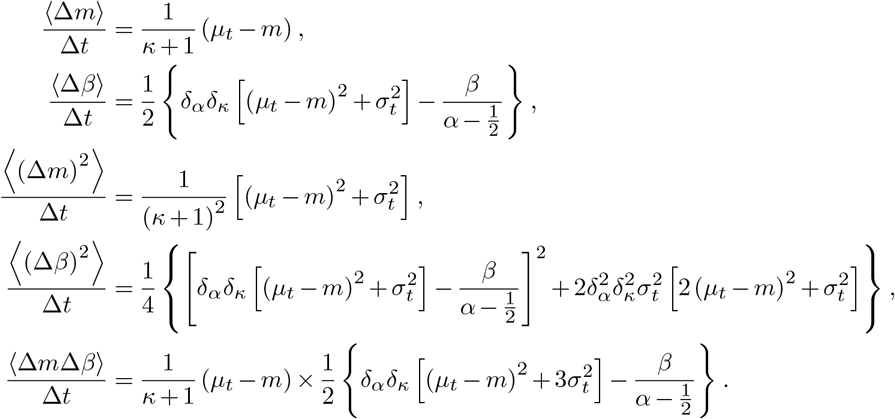

#### Fokker-Planck equation

Equation (8) is a stochastic differential equation that describes the evolution of ***θ*** = (*m, β*) for a single immune cell. To understand the distribution of *m* and *β* at the population level, we derive the Fokker-Planck equation corresponding to equation (8). The Fokker-Planck equation describes the evolution of the probability distribution over ***θ*** parameters in a population where the dynamics of each individual is governed by equation (8). Following standard methods ^56^, the form of the Fokker-Planck equation is

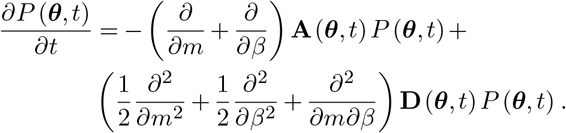

Substituting in the expressions for **A** and **D** derived above, we obtain

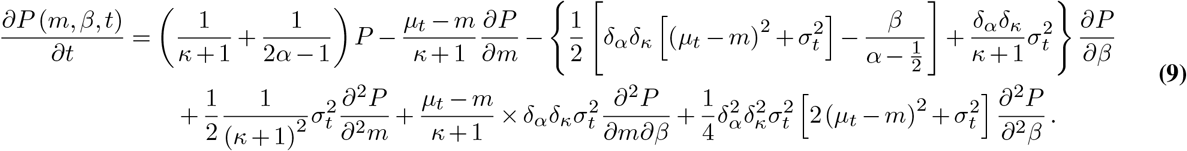

#### Numerical integration of the Fokker-Planck equation

We used the central difference method to solve equation (9) numerically. Here we simulated adaptation to a true signal distribution with mean *μ_t_* = 2 and standard deviation *σ_t_* = 1, using the memory parameters *κ* = 20 and *α* = 10. We anticipated that the learned signal parameters should be centered around *m* = 2 and *β* = 9, which would give 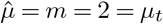 and 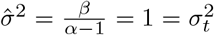. Thus we selected a rectangular domain [1.5, 2.5] × [5, 15] in the (*m, β*) space to numerically evaluate the equations (see **Fig. 4**). We set *h* = *w* = 0.01 as the spatial discretization size of *m* and *β*, respectively, dividing the domain into a 100 × 1000 grid. The discrete first and second derivatives at each grid point can be expressed as

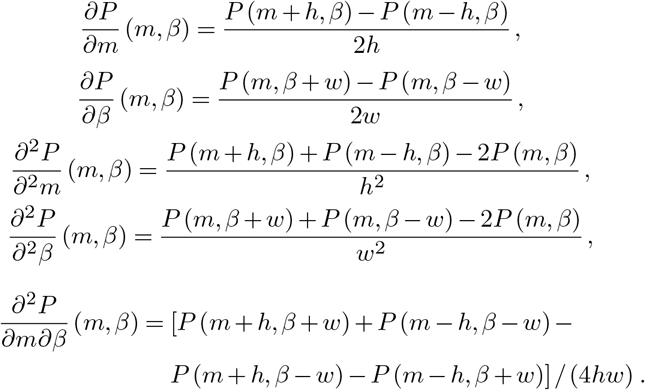

After applying the spatial discretization to the right hand side, equation (9) can be simplified to

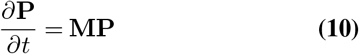

where the discrete probability distribution **P** is a 10^5^ dimensional vector representing values of the function at all grid points. **M** is a 10^5^ × 10^5^ matrix with entries determined by equation (9), using the discrete derivative formulas above. **M** does not change over time.

We used the Crank–Nicolson method ^57^, which is numerically stable and second-order in time, to solve equation (10). The Crank–Nicolson discretization is

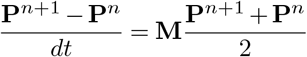

where *n* is the discrete time index, proportional to the number of interactions between immune cells and target cells. This is a linear equation with variables **P**^*n*+1^ and **P**^*n*^. It can be written in the form of 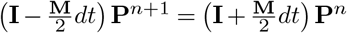, where **P**^*n*+1^ is solved for and normalized to 1 at each step. The initial values of the probability distribution *P* (*m, β,* 0) were set to be uniform among the internal grid points and zero at the boundaries. The time step was set as *dt* = 0.001.

## Data and code

Data and code used in our analysis is available at the GitHub repository https://github.com/bartonlab/paper-innate-immune-adaptation. This repository also contains Jupyter notebooks that can be run to reproduce the results presented here.

---

**Supplementary Fig. 1.**
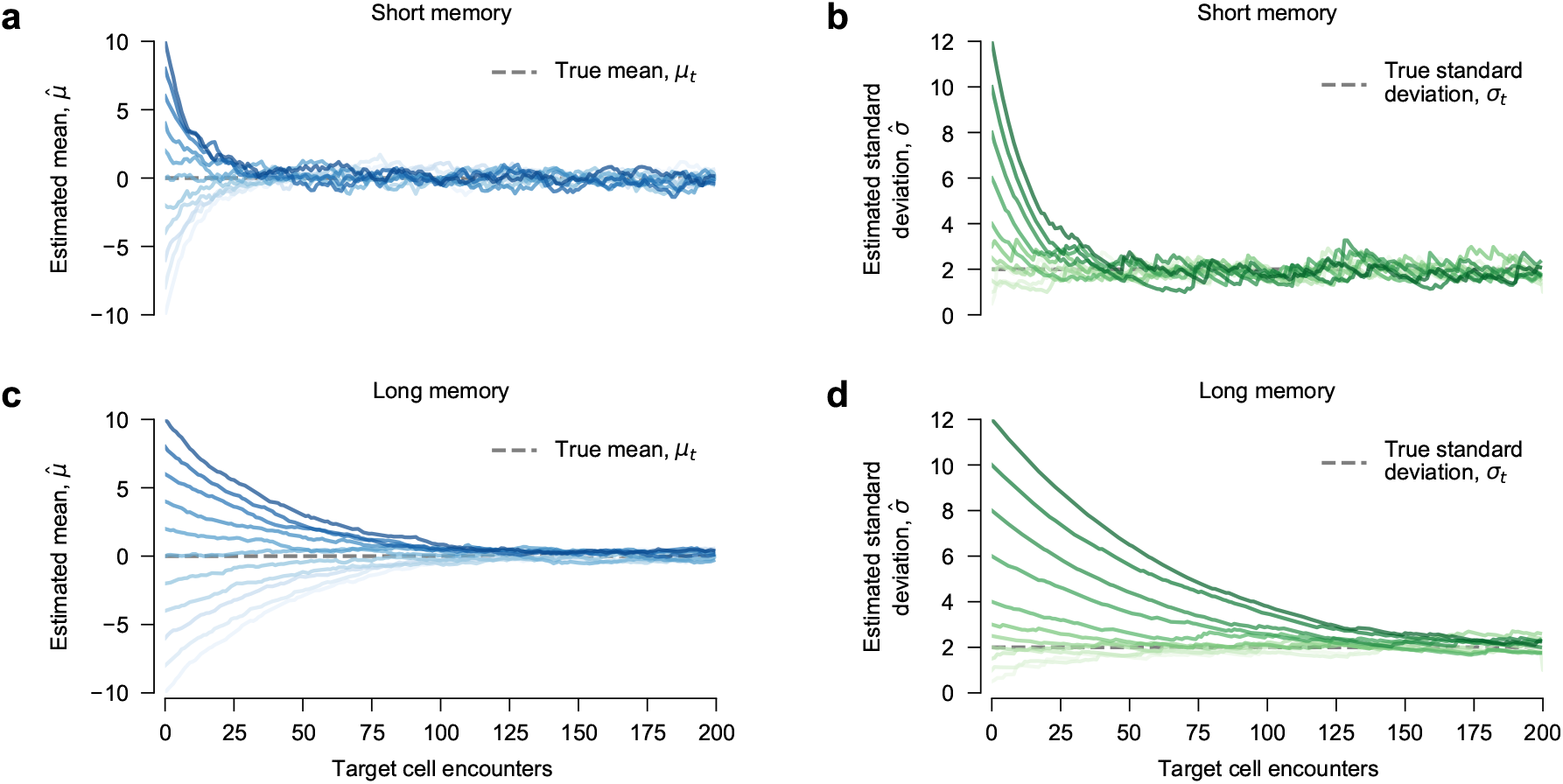
Immune cells with shorter memory lengths exhibit faster but noisier adaptation. The estimated mean 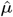 and standard deviation 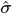 approach the true mean (*μ_t_* = 0) and true standard deviation (*σ_t_* = 2) in a finite number of encounters, regardless of the initial value and memory length. When the memory length is shorter (*κ* = 10 and *α* = 5, panels **a** and **b**), estimated values approach the true ones faster than for immune cells with longer memory lengths (*κ* = 40 and *α* = 20, panels **c** and **d**). However, adaptation is less precise when the memory length is shorter. **a**, Convergence to true mean (*μ_t_* = 0) from various initial values of *m*. The initial value of *β* = (*α* − 1) is the same in all cases. **b**, Convergence to the true standard deviation (*σ_t_* = 2) from various initial values of *β*. The initial value of *m* = 0 is the same in all cases. The memory length for both **a** and **b** is set by *κ* = 10 and *α* = 5. (**c**, **d**) Display convergence toward the true mean and standard deviation as in (**a**, **b**). Initial parameters are the same as those in (**a**, **b**), except for longer memory lengths *κ* = 40 and *α* = 20.

**Supplementary Fig. 2.**
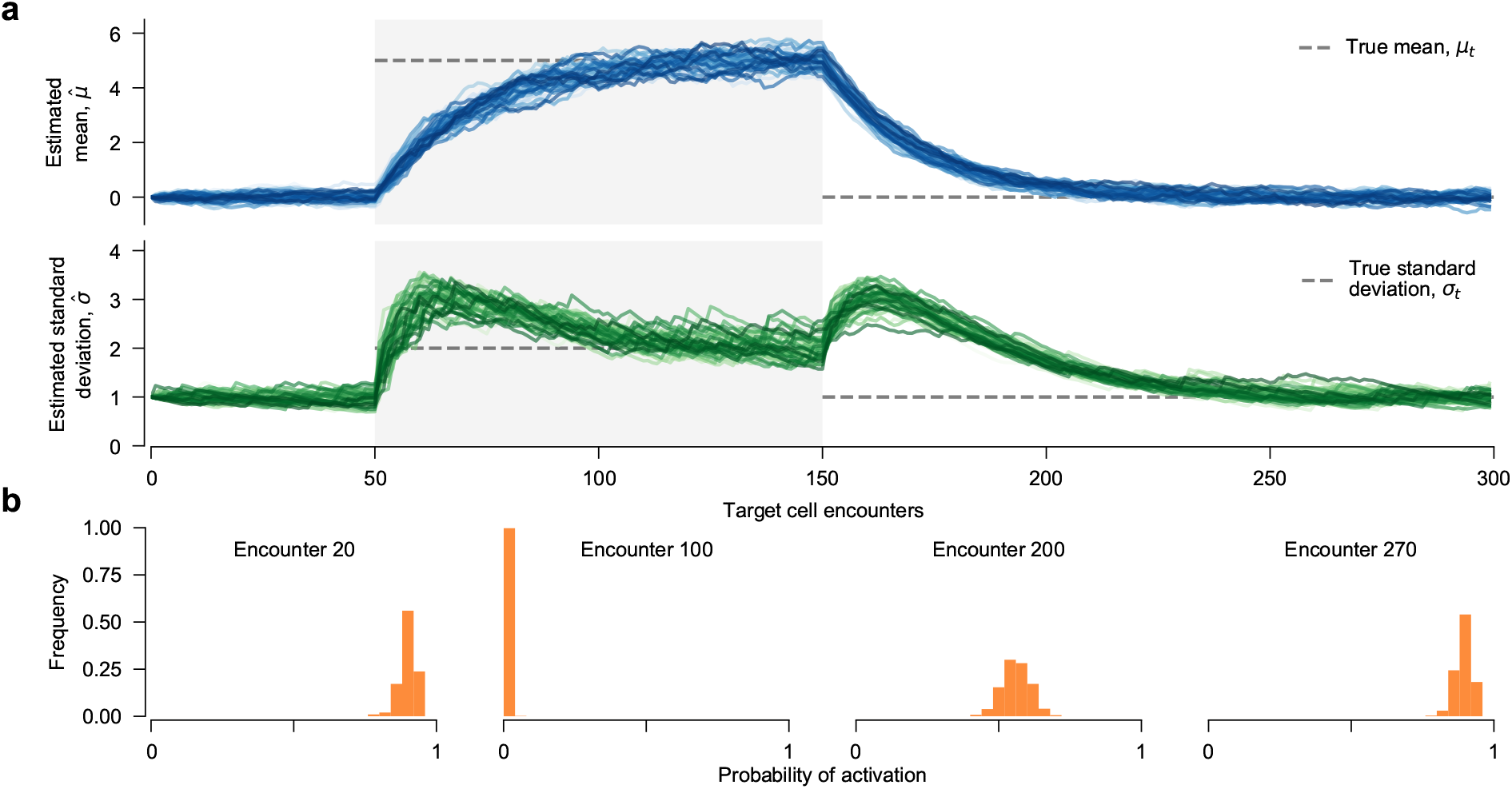
Finite memory of past interactions with target cells results in heterogeneous immune cell behaviors. **a**, Adaptation of 500 immune cells to changing environments, following the same conventions as in Fig. 3. Even though all immune cells start with the same initial values (*m* = 0 and *β* = 9, with memory parameters *κ* = 20 and *α* = 10), the learned distribution for each immune cell evolves differently over time. This is due to the stochastic nature of signals from target cells, and the finite memory length of past target encounters. **b**, The heterogeneity of learned signal distributions also leads to heterogeneous immune cell responses, characterized by the probability of activation against an aberrant target cell. The signal distribution for aberrant targets is the same as in Fig. 3.

**Supplementary Fig. 3.**
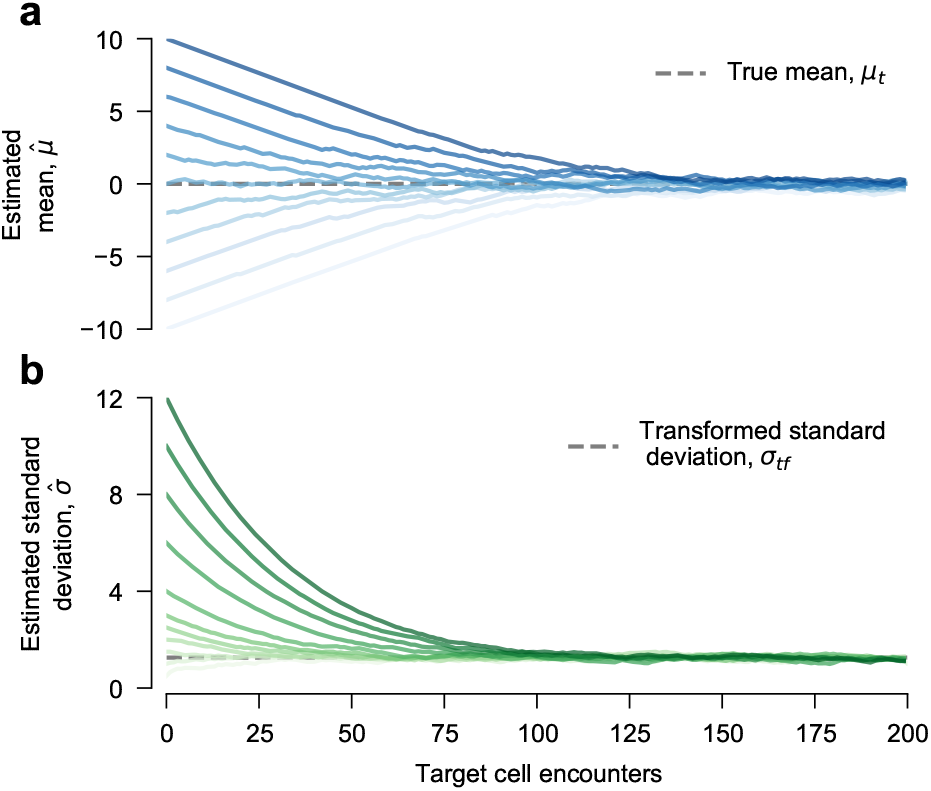
Immune cell adaptation is more gradual with signal saturation. The estimated mean. (**a**) and standard deviation (**b**) converge toward the true mean and transformed standard deviation in a model with signal saturation (see Methods). Parameters and initial conditions are the same as those in Fig. 2. Signal saturation changes the expected standard deviation of the signal when the model is perfectly adapted because large deviations are suppressed. We have thus replaced the true standard deviation *σ_t_* with the transformed standard deviation *σ_tf_* in **b** (Methods).

**Supplementary Fig. 4.**
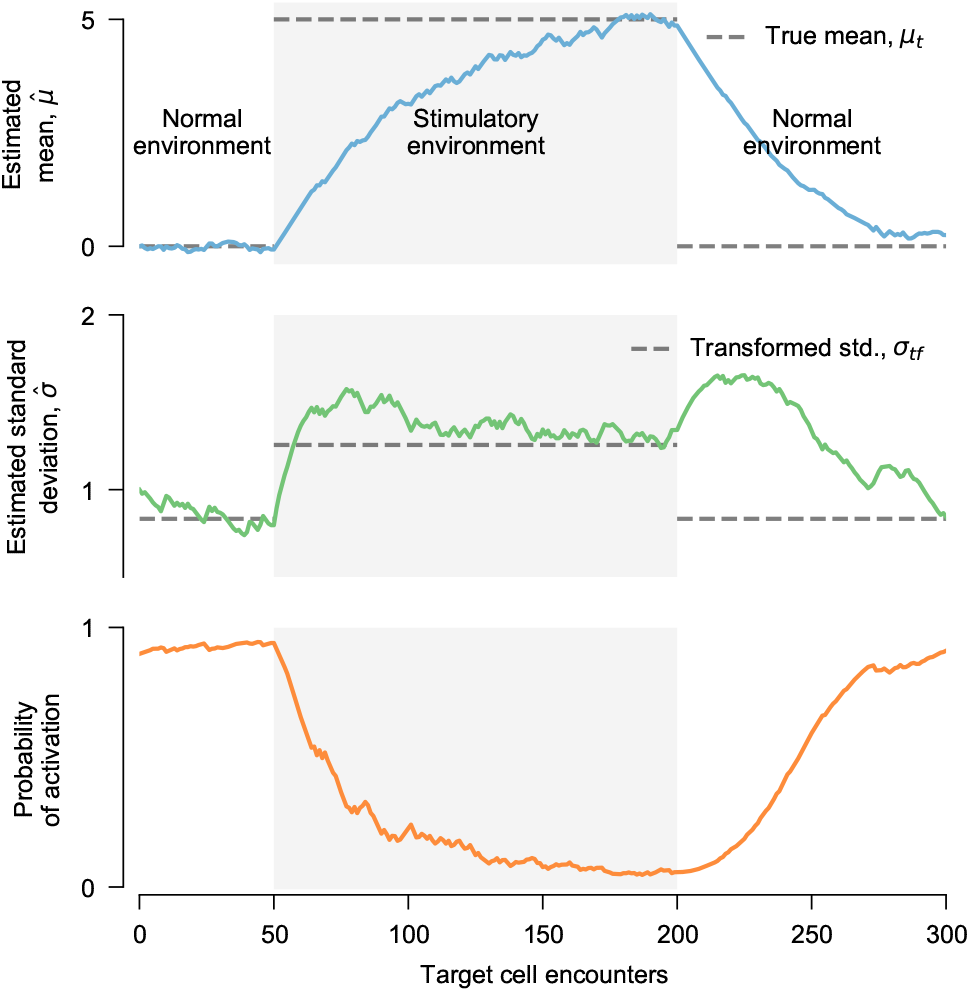
Immune cell adaptation is more gradual with signal saturation. As in Fig. 3, an immune cells adapts to different signal distributions in ‘normal’ and ‘stimulatory’ environments in a model with signal saturation (see Methods). Adaptation is more gradual due to signal saturation, so we have extended the number of encounters in the stimulatory environment to allow for complete adaptation. As in Fig. 3, the immune cell is able to reliably activate against aberrant targets when it is adapted to a normal environment, but it loses this ability after long times in a stimulatory environment. Compared to a model without signal saturation (Fig. 3), adaptation is more gradual and less noisy. In addition, the immune cell retains the ability to activate against aberrant targets for a longer time after being placed in the stimulatory environment.

